# Generalized tree structure to annotate untargeted metabolomics and stable isotope tracing data

**DOI:** 10.1101/2023.01.04.522722

**Authors:** Shuzhao Li, Shujian Zheng

## Abstract

In untargeted metabolomics, multiple ions are often measured for each original metabolite, including isotopic forms and in-source modifications, such as adducts and fragments. Without prior knowledge of the chemical identity or formula, computational organization and interpretation of these ions is challenging, which is the deficit of previous software tools that perform the task using network algorithms. We propose here a generalized tree structure to annotate ions to relationships to the original compound and infer neutral mass. An algorithm is presented to convert mass distance networks to this tree structure with high fidelity. This method is useful for both regular untargeted metabolomics and stable isotope tracing experiments. It is implemented as a Python package (khipu), and provides a JSON format for easy data exchange and software interoperability. By generalized pre-annotation, khipu makes it feasible to connect metabolomics data with common data science tools, and supports flexible experimental designs.

## Introduction

Metabolomics is becoming an increasingly important tool to biomedicine. Untargeted LC-MS (liquid chromatography-mass spectrometry) metabolomics is key to perform high-coverage chemical analysis and discoveries. The term “annotation” in metabolomics often includes i) the assignment of measured ions to their original compounds, and ii) establishing the identity of the compounds (Domingo-Almenara et al, 2018; Blaženović et al, 2019). For clarity, we refer the first step as “pre-annotation” in this paper, which is the assignment of isotopes, adducts and fragments to the unique compounds. Correct pre-annotation will greatly facilitate the later step of identification, by reducing errors on analyzing and searching the redundant ions. Multiple software tools have been developed for this purpose of pre-annotation, including CAMERA (Kuhl et al, 2012), Mz.unity (Mahieu et al, 2016), xMSannotator (Uppal et al, 2017), MS-FLO (DeFelice et al, 2017), MetNet (Naake and Fernie, 2018), CliqueMS (Senan et al, 2019), Binner (Kachman et al, 2020) and NetID (Chen et al, 2021).

In high-resolution mass spectrometry, the *m/z* (mass to charge ratio) difference between isotopes is usually resolved unambiguously. Adducts are formed in the ionization process, therefore, those from the same original compound should have the same retention time in chromatography. Besides adduct ions, formation of conjugates and fragments (including neutral loss) also belongs to in-source modifications. Isotopes, adducts and fragments are often referred as redundant or degenerate peaks in LC-MS literature. All pre-annotation tools utilize the *m/z* differences between peaks, which correspond to the mass differential between isotopes, or between atoms or chemical groups. In addition, having the same retention time is a critical requirement to group these degenerate peaks. Some tools also use similarity in the shape of elution peaks and sometimes statistical correlation between peak intensity across samples. Such correlations can be supporting evidence but are not a prerequisite (Mahieu et al, 2016).

Most pre-annotation tools use a network representation of degenerate peaks. Because the pairwise relationships between peaks are established first, then it is natural to connect the pairs into networks by using pairwise relationships as edges and shared peaks as nodes. Such networks still contain redundant and often erroneous edges. The main challenge remains to resolve how all peaks are generated from the same original compound, which requires a) inferring the neutral mass of the original compound, and b) establishing the relationship of all peaks to the original compound. Given the difficulty of organizing this information in untargeted metabolomics, the coverage of untargeted analyses is often called to question.

A couple of notable studies tried to address the question of coverage using isotope tracing in untargeted metabolomics, and suggested that a small number of metabolites are actually measured and the majority of peaks are “junks", either from contaminations, isotopes or LC-MS artifacts (Mahieu and Patti, 2017; Wang et al, 2019). A new challenge also arose that analyzing these isotope tracing data by global metabolomic is not trivial. So far, isotope tracing experiments usually require targeted metabolites and specialized software (Chokkathukalam et al, 2013; Bueschl et al, 2017; Previs and Downes, 2020; Rahim et al, 2022). In untargeted analysis, without prior knowledge of the chemical formulas, special experimental designs are required and the software tools are tied to the designs, which are the cases for X^13^CMS (Huang et al, 2014; Llufrio et al, 2019) and PAVE (Wang et al, 2019). It is highly desirable to have a generic and flexible tool to process untargeted isotope tracing metabolomics, and to enable more flexible data analysis and modeling.

In this study, we propose a generalized tree structure to assign relationship of each ion to the original compound and infer its neutral mass. The pre-annotation software tool, khipu, is freely available as a Python package. It is applicable to both regular untargeted metabolomics and stable isotope tracing data, and helps plug metabolomics data easily into common data science tools.

## Results

### The combination of isotopes and adducts is a 2-tier tree

The redundant or degenerate ions in mass spectrometry can be from in-source modifications (adducts, fragments and conjugates) on any of the isotopic forms. For simplicity, we only consider adducts in the initial steps. The combination of isotopes and adducts leads to a grid of mass values, relative to the neutral mass of M0, exemplified in **Table 1**. We use M0 to denote the molecules with only 12C atoms. The isotopes are denoted as 13C/12C, 13C/12C*2, etc., whereas the last digit is the number of 13C atoms present in each molecule.

**Table 1.** Combinations of isotopes and adducts generate mass differences as a grid. The mass values are relative to the 12C only neutral mass. Examples are using a limited number of isotopes and in-source modifications in positive ionization.

|  | *M+H[+]* | *M+NH4[+]* | *M+Na[+]* | *M+HCl+H[+]* | *M+K[+]* | *M+ACN+H[+]* |
| --- | --- | --- | --- | --- | --- | --- |
| *M0* | 1.007276 | 18.033826 | 22.989276 | 36.983976 | 38.963158 | 42.033825 |
| *13C/12C* | 2.010631 | 19.037181 | 23.992631 | 37.987331 | 39.966513 | 43.03718 |
| *13C/12C\*2* | 3.013986 | 20.040536 | 24.995986 | 38.990686 | 40.969868 | 44.040535 |
| *13C/12C\*3* | 4.017341 | 21.043891 | 25.999341 | 39.994041 | 41.973223 | 45.04389 |
| *13C/12C\*4* | 5.020696 | 22.047246 | 27.002696 | 40.997396 | 42.976578 | 46.047245 |
| *13C/12C\*5* | 6.024051 | 23.050601 | 28.006051 | 42.000751 | 43.979933 | 47.0506 |
| *13C/12C\*6* | 7.027406 | 24.053956 | 29.009406 | 43.004106 | 44.983288 | 48.053955 |

The adducts can be represented as a tree (**Figure 1A**), using the neutral form as the root, which is usually not measured in mass spectrometry. Each edge in the tree corresponds to a specific mass difference, from the reaction forming the adduct. In fact, the full grid in Table 1 can be accommodated into the tree, using isotopes as leaves to the adducts. Two arguments favor the tree as a preferred data structure over a generic network: 1) each ion measured in mass spectrometry is formed from a specific “predecessor”, and 2) the whole group of ions are from a unique compound, which is the “root”. In computational terms, a network becomes a tree once it fulfills the two requirements: 1) each node can have no more than one predecessor and 2) a unique root. The benefit of this tree representation is important, allowing automated interpretation of all ions via defined semantics.

**Figure 1.**
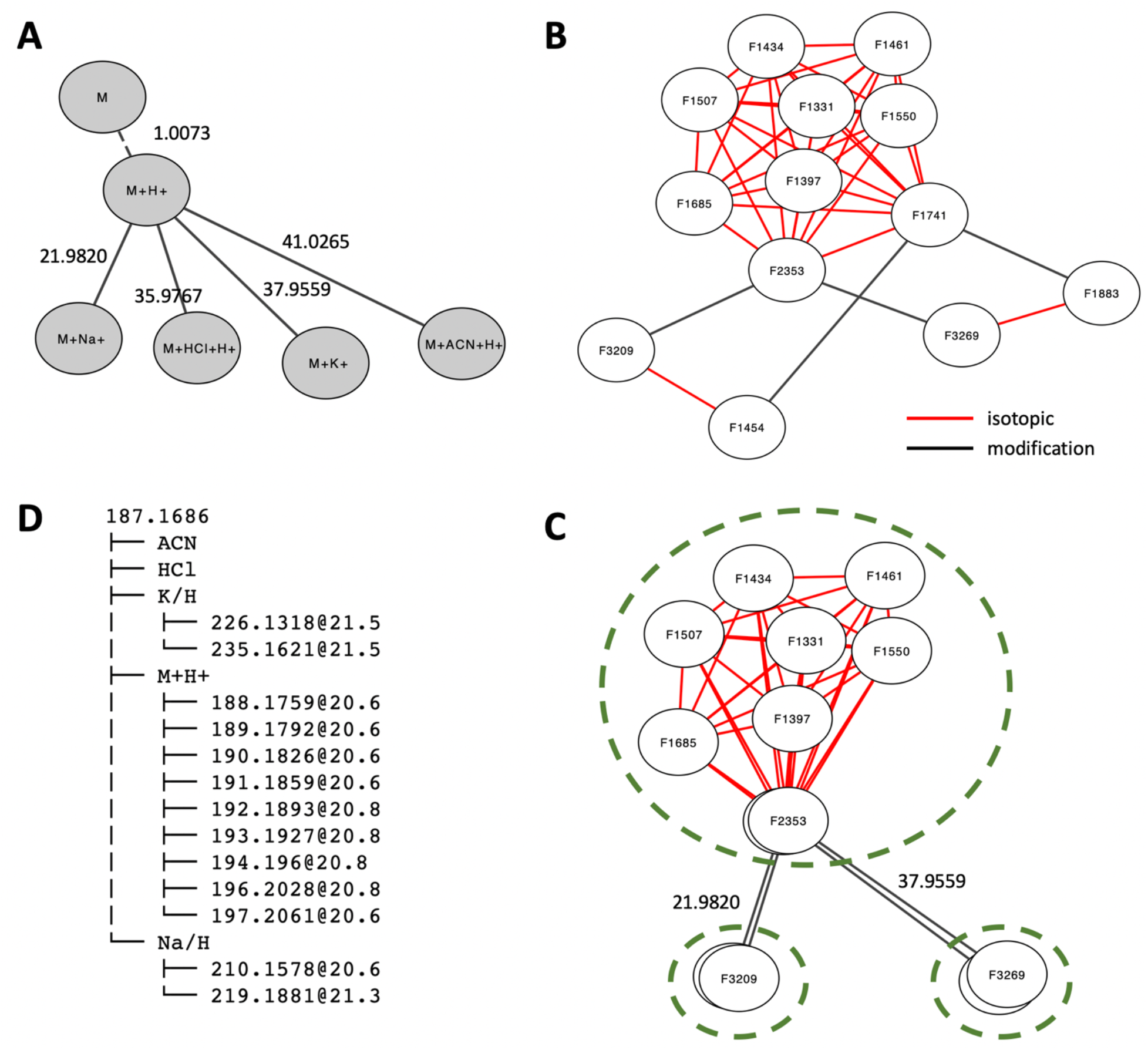
The khipu algorithm converts a mass distance network to a tree structure. A) An adduct tree base on Table 1. Mass differences on the edges are relative to the predecessor nodes. B) An example mass distance network from our credentialed E. coli dataset, which contains both unlabeled and 13C labeled samples. Edges in red are from isotopic patterns and edges in black from adduct patterns. C)The isotopic subnetworks can be treated as individual nodes, then the abstracted network has only adduct edges, which facilitates the alignment to the theoretical adduct tree in A). D) Resulted 2-tier tree. The root is inferred neutral mass. No ion is assigned to ACN or HCl adducts. Decimal numbers should be consistent with that in Table 1.

Because the isotopes are present independently from each other at the time of measurement, we treat them equally as one tier of the tree here. It is noted that the generation of them may have biochemical significances in isotope tracing experiments, but that problem is outside data processing and annotation. Therefore, the combination of isotopes and adducts, as exemplified in Table 1, can be represented as a 2-tier tree. The tree can either use adducts as tier 1 or isotopes as tier 1. The decision is to use adducts as tier 1, because a) adduct mass patterns are more distinct, and b) isotopes are often limited by abundance, resulting only M0 ions in many compounds.

### An algorithm to convert a mass distance network to a 2-tier tree

Annotation methods in MS metabolomics commonly start by searching mass difference patterns, e.g. 1.0034 for 13C/12C in isotopes and 22.9893 for Na^+^ in adducts. Each match leads to a pair of ions (also called features), and many pairs are connected via shared ions into a network of ions (**Figure 1B**). During the mass difference search, additional redundancy is introduced, e.g. the mass difference between 13C and 12C is the same as between 13C/12C*2 and 13C/12C*3, and so forth. This network redundancy is apparent in the top part of the network in Figure 1B. The objective in annotation is to identify the true root (original compound) from the network, which has been challenging in previous works.

As biological reactions are not part of data annotation here, the edges in our mass distance networks belong to one of the two categories: isotopic differences or in-source modifications (**Figure 1B**). A key observation is that all ions connected by isotopic edges belong to the same adduct. Therefore, subnetworks per adduct can be defined from a mass distance network (**Figure 1C**). Once these isotopic subnetworks are abstracted into individual network nodes, we can find the best alignment between this abstracted network (**Figure 1C**) and the adduct tree (**Figure 1A**). The algorithm is designed as two-step optimization: to obtain a tree with optimal number of ions explained in the alignment of adduct trees, then in the alignment of isotopes. The result of this algorithm on our example network is shown in **Figure 1D**. To match our 2-tier tree structure, the networks have to become directed acyclic graph (DAG). During this process, erroneous edges are weeded out because they do not satisfy DAG and a rooted tree. This method yields a structured and unique annotation of each ion in the tree. Based on the matched *m/z* values, the neutral mass of M0 compound is obtained by a regression model. Once the core structure of a tree is established, additional adducts and fragments can be searched in the data. The algorithm is implemented into a freely available Python package khipu.

### Khipu plots allow intuitive interpretation of isotope tracing data

After ions are grouped into a tree for each original compound, they are recorded into transparent JSON format, as defined for empirical compounds (see examples in **Supplemental notebook**). An “empirical compound” refers to a tentatively defined compound in metabolomics data, used in our previous projects (Li et al, 2013, Pang et al, 2020), as the technology may not deliver definitive identification or resolve a mixture (e.g. isomers not successfully separated).

We continue using the compound in **Figure 1 B-D** to illustrate the khipu plotting functions. Each ion is measured with an intensity value in one of more biological samples. While the tree visualization in Figure 1D is useful, khipu includes multiple functions to visualize the features, *m/z* values, intensity values as data frame tables (Supplemental notebook), to facilitate intuitive interpretation of each compound. An enhance visualization of the tree is demonstrated in **Figure 2A**, where the adducts are organized as a “trunk” and isotopes as “branches”. It’s clear that several isotopes are present as the protonated ion; Na and K adducts are present for the more abundant isotopes. Because this visualization style resembles the khipu knot records used by Andean South Americans, we named our software “khipu”.

**Figure 2.**
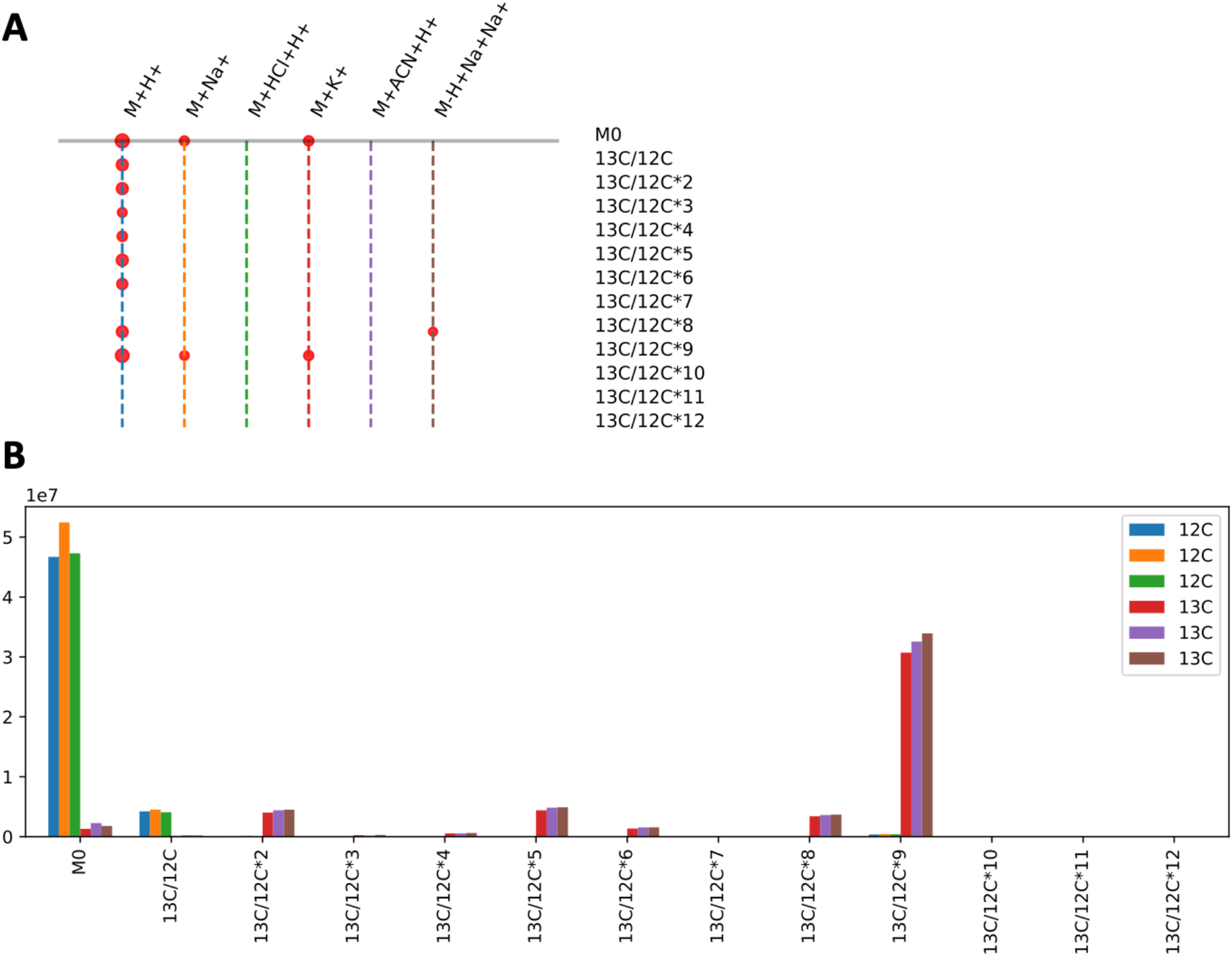
Visualization using khipu facilitates interpretation of isotope tracing data. A) An example khipugram plot for the compound in Figure 1, with its 13 ions aligned to the tree in Figure 1D. Each dot represents an ion measured in the data, the size of dots proportional to average intensity. The vertical dashed lines are colored for easy navigation, and the colors are of no particular meaning. B) Bar plot for intensity values of the M+H^+^ ion in different isotopes (x-axis) for three 12C samples and three 13C samples (in color legend). This is from the first branch in A).

This experimental dataset was from cultured E. coli, containing three unlabeled samples and three samples grown on U-13C-glucose. **Figure 2B** visualizes the intensity values across samples for the M+H^+^ ion. The three unlabeled samples have high M0 peaks, and smaller 13C/12C (M1) peaks due to the naturally occurring isotopes. The U-13C labelled samples have the highest peaks at 13C/12C*9 (M9), with smaller peaks of other isotopes. This indicates that the latter samples are almost fully labelled by 13C, and the compound should contain 9 carbon atoms. The neutral mass inferred by khipu is 187.1686, which matches to acetylspermidine, which has a chemical formula C9H21N3O, perfectly consistent with the isotopic pattern.

As a pre-annotation took, khipu is positioned to feed organized data for downstream data analysis. Users can choose to model the isotopes and compute flux using other tools (Moseley 2010; Millard et al, 2012). Khipu results can be easily used by other software and analyzed using common data science tools (demo notebooks included in the code repository). JSON (JavaScript Object Notation) is a common format for data exchange between software programs and web applications, and one of khipu export formats. This enables an effective way for sharing metabolite annotation, which is human friendly, computable, and neutral to software platforms. A snippet of khipu export in JSON is as follows:

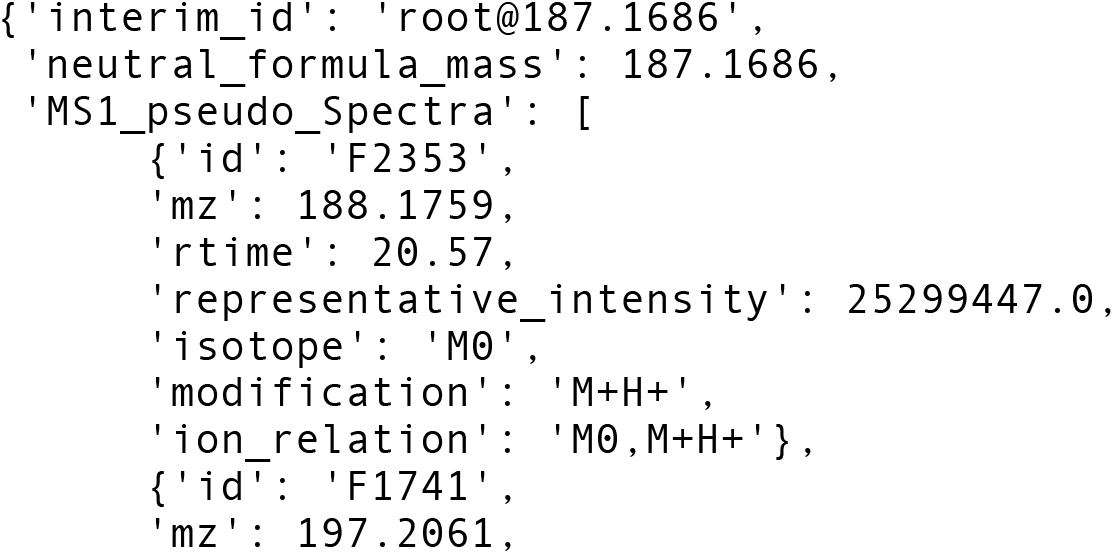
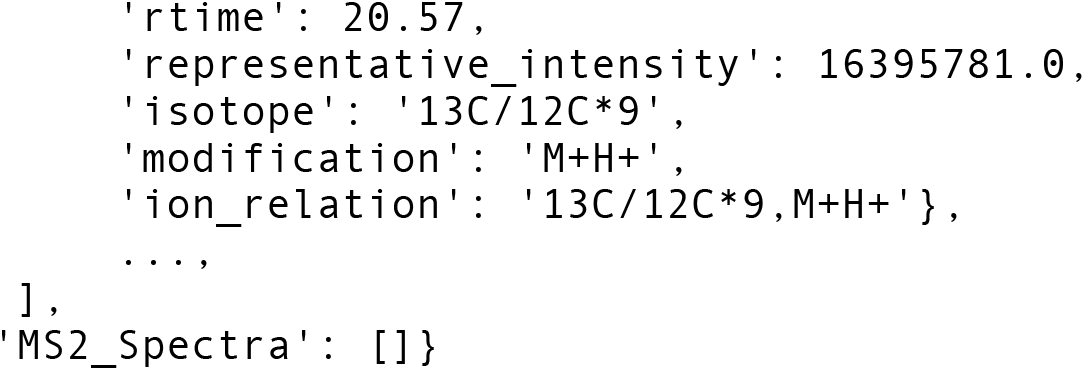

### How many metabolites do we measure?

Proper pre-annotation is key to answer the question of how many metabolites/compounds are measured in an experiment, which is a matter that has been debated for over a decade. Many studies overestimated the coverage because the database search was inflated by redundant/degenerate features/ions. Studies from the Patti and Rabinowitz labs used isotope tracing techniques, and suggested the numbers are around 1,000~2,000 in E. coli and yeast (Mathieu and Patti, 2017; Wang et al, 2018). Our khipu software now provides systematic and fast pre-annotation on metabolomic datasets.

In our E. coli data (reverse phase ESI+), 3,602 LC-MS features were measured, and khipu annotated 548 empirical compounds (trees) from 1,745 features. Among the 548 empirical compounds, 445 have multiple isotopes (**Figure 3A**). The remaining 1,857 features are singletons, i.e., not grouped with any other features. In two yeast datasets from Rabinowitz lab, khipu annotation resulted in 1,775 and 908 empirical compounds, respectively in ESI+ and ESI-modes (**Figure 3B&C**). In the yeast datasets, we included additional adducts from Wang et al (2021), which by design did not increase the number of empirical compounds, but increased the explained ESI+ features from 6,310 to 8,049, and from ESI-features 2,601 to 2,912. These results suggest that less than 2,000 compounds were reliably measured in these experiments. Of note, closer examination of each dataset should also remove contaminants, which is not part of khipu.

**Figure 3.**
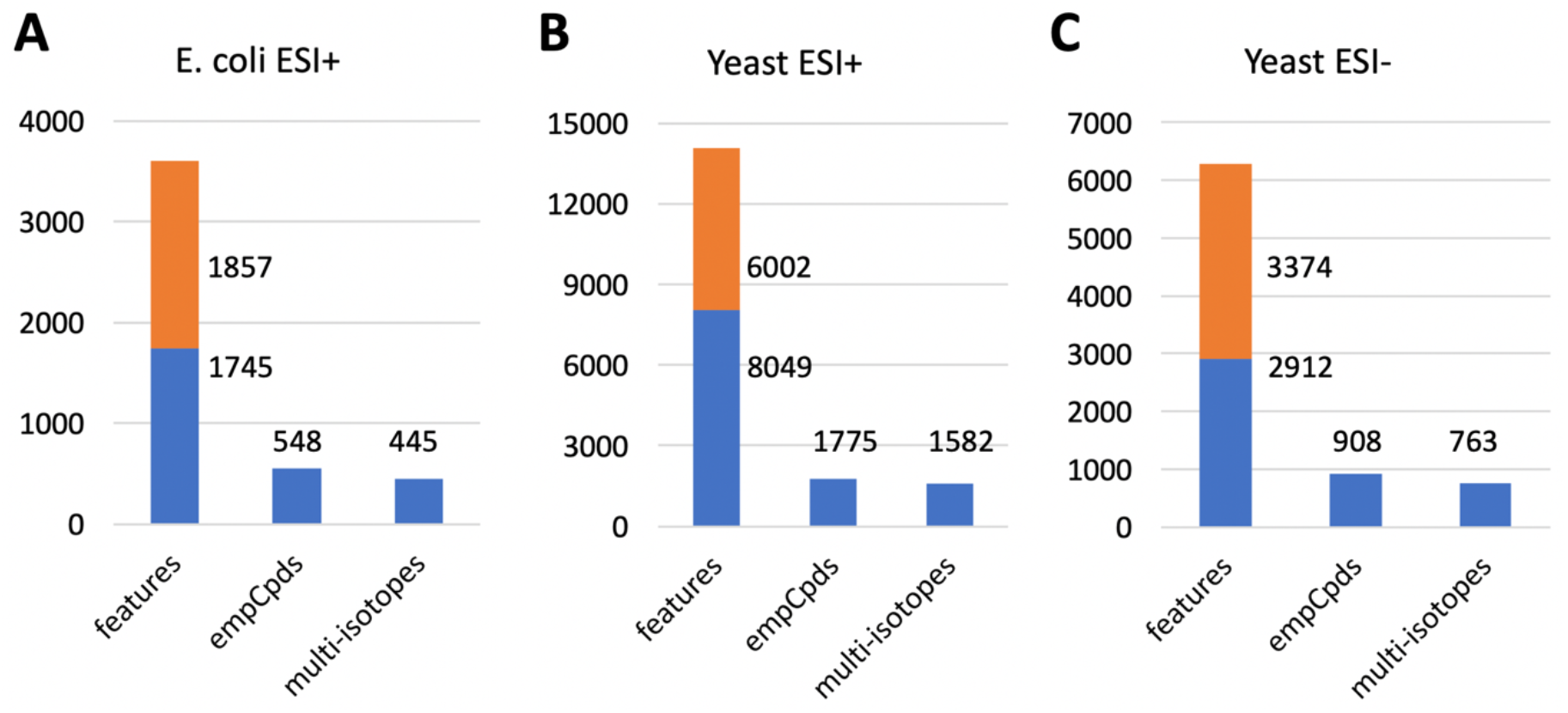
Number of measured compounds in three metabolomic datasets. A) Credentialed E. coli data generated in this study. B) Previously published yeast ESI+ and C) ESI-datasets from Rabinowitz lab (Wang et al. 2021). Khipu annotation on these datasets took 2~6 seconds on a laptop computer of Intel i7 CPU. The orange portions are referred as “singletons”.

## Discussion

Annotation of untargeted metabolomics data, including isotopic tracing data, is still not fully solved. Many current tools take a network approach but depend on assumed base ions or formulas to assign relationship between ions. We present a new algorithm here to resolve the mass distance networks into a tree structure, unambiguously defining ion relationships and inferring neutral mass. This approach shall reduce false annotations, and facilitate new compound identification and discoveries. We consider the pre-annotation with khipu a key step forward, also because it ships with generalized annotation format, which will greatly facilitate data exchange and software interoperability. With this foundation, future benchmarking and improvements are expected.

Multiple Jupyter notebooks are provided as part of the software package to demonstrate how khipu is plugged into common data science tools. This gives great flexibility to people in using both regular and isotope tracing metabolomics data, because the computational methods, as well as experimental designs, are no longer limited by rigid software designs. This is an emerging model in data science. Traditional software development is often too costly, and its maintenance is even more challenging (Chang et al, 2021). Fundamentally, no software developer can meet every demand via point-and-click interface. Scientific data analysis has to depend greatly on scripting. The combination of modular software components, transparent data structures and Jupyter notebooks opens up many opportunities for collaborations and scientific progress (Pittard et al, 2020).

Khipu can be easily reused by other software tools. We plan to integrate it with the preprocessing software asari (Li et al, 2022), whereas elution patterns can be better determined than from standalone feature tables. The standard input to khipu is tab delimited feature tables, which should be compatible with any LC-MS preprocessing software. Therefore, it will be easy to incorporate it into metabolomics workflows, where complete annotation can take into consideration of contaminants, authentic libraries and tandem mass spectrometry data.

## Methods

### Python implementation

Khipu is developed as an open source Python 3 package, and available to install from the standard PyPi repository via the pip tool. It is freely available on GitHub (https://github.com/shuzhao-li/khipu) under a BSD 3-Clause License. The graph operations are supported by the networkx library, tree visualization aided by the treelib library. Khipu uses our package mass2chem for search functions. The data model of “empirical compound” is described in the metDataModel package. The package is designed in a modular way to encourage reuse. The classes of Weavor and Khipu contain main algorithms, supported by numerous utility functions. All functions are documented in the source via docstrings. Examples of reuse are given in wrapper functions and in Jupyter notebooks. It can be run as a standalone command line tool. Users can use a feature table from any preprocessing tool as input and get annotated empirical compounds in JSON and tab delimited formats.

### LC-MS metabolomics data

The dry extracts of unlabeled and ^13^C labeled *E. coli* (Cambridge Isotope Laboratories, Inc.; Catalog number: MSK-CRED-DD-KIT) were reconstituted in 100 μL of ACN/H_2_O (1:1, v/v) then sonicated (10 mins) and centrifuged (10 mins at 13,000 rpm and 4°C) before overnight incubation at 4°C. The supernatant for each ^12^C/^13^C *E. coli* extract was collected and then prepared for LC-MS analysis. Metabolite extraction was carried out using acetonitrile:methanol (8:1, v/v) containing 0.1% formic acid. All samples were vortexed and incubated with shaking at 1000 rpm for 10 min at 4°C followed by centrifugation at 4°C for 15 min at 15,000 rpm. The supernatant was transferred into mass spec vials and 2 μl injected into UHPLC-MS. All samples were maintained at 4 °C in the autosampler, and analyzed using a Thermo Scientific Orbitrap ID-X Tribid Mass Spectrometer coupled to a Thermo Scientific Transcen LX-2 Duo UHPLC system, with a HESI ionization source, using positive ionization. A Hypersil GOLDTM RP column (3 μm, 2.1 mm x 50 mm) maintained at 45 °C was used. 0.1% formic acid in water and 0.1% formic acid in acetonitrile were used as mobile phase A and B respectively. The following gradient was applied at a flow rate of 0.4 ml/min: 0-0.1 min: 0% B, 0.10-1.9 min: 60% B, 1.9-5.0 min: 98% B, 5.00-5.10 min: 0% B and 4.9 min cleaning and column equilibration. The chromatographic run time was 5 min followed by 5 min washing step after each sample. The MS settings are: spray voltage, 3500 V; sheath gas, 45 Arb; auxiliary gas, 20 Arb; sweep gas, 1 Arb; ion transfer tube temperature, 325 °C; vaporizer temperature, 325 °C; mass range, 80-1000 Da; maximum injection time, 100 ms; resolution 60,000.

The yeast data from Chen et al. 2021 were retrieved from the MassIVE repository (https://massive.ucsd.edu, ID no. MSV000087434). The yeast ESI+ data contain both unlabeled and 13C isotope labeled samples, while the ESI-data did not involve isotope tracing, The data from Mathieu and Patti (2017) and Wang et al (2018) were not found publicly. All datasets were processed using asari version 1.9.2 (https://github.com/shuzhao-li/asari). The yeast ESI-dataset was quality filtered for signal-noise-ratio > 100 to serve as a cleaner demo.

## Supplemental File

**data_analysis_ecoli_pos.pdf**

A Jupyter Notebook printed to PDF format to demonstrate khipu applications.

## Code availability

The asari source code is available at GitHub, https://github.com/shuzhao-li/khipu, and as a Python package via https://pypi.org/project/khipu-metabolomics/. The demonstration datasets are provided as part of the source code.

## Acknowledgements

This work was in parted funded by NIH grants (to SL) U01 CA235493 (NCI) and R01 AI149746 (NIAID).

## Author contributions

S.L. designed the study, wrote the khipu software and the manuscript. S.Z. performed the LC-MS metabolomics experiment on credentialed E. coli samples.

