## Supplemental File 1 for "Generalized tree structure to annotate untargeted metabolomics and stable isotope tracing data"

### Khipu demo for ecoli_pos data analysis

This notebook is run on $docker run -v /Users/shuzhao/play:/home/jovyan -p 8888:8888 jupyter/scipy-notebook

#### Part I. Differential features between 12C and 13C samples

The data contain 6 samples, 3 in each group. We first do a t-test to get features that differ between two groups.

In [1]:
```
import pandas as pd
import numpy as np
import matplotlib.pyplot as plt
from scipy.stats import ttest_ind
```

In [2]:
```
# demo data are at https://github.com/shuzhao-li/khipu/tree/main/testdata/
# Have a look via pd
ecoli = pd.read_table("ecoli_pos.tsv", index_col=0, sep="\t")
ecoli.head()
```

Out[2]:

| | mz | rtime | 12C_Ecoli_20220321_004 | 12C_Ecoli_20220321_004_20220322095030 | 12C_Ecoli_20220321_004_20220322130235 | 13C_Ecoli_20220321_0 |
|---|---|---|---|---|---|---|
| id_number | | | | | | |
| F1 | 61.9278 | 40.94 | 42543 | 0 | 0 | 433 |
| F2 | 95.9735 | 125.06 | 2079328 | 984349 | 1314348 | 17078 |
| F3 | 115.9802 | 11.42 | 3362695 | 3228128 | 4076328 | 32017 |
| F4 | 115.9860 | 5.83 | 2035714 | 1739145 | 1978764 | 19778 |
| F5 | 115.9860 | 27.96 | 182603 | 219669 | 248607 | 1990 |

In [3]:
```
# t-test of each row between 12C and 13C samples
# data are log2 transformed in this test to have more normal distribution; +1 to avoid log2(0)
def ttest(row):
    t,p = ttest_ind(np.log2(row[3:6]+1), np.log2(row[6:9]+1))
    return p

pvalues_featurelist = ecoli.apply(ttest, axis=1)
```

In [4]:
```
# sort p-values
pvalues_featurelist = pvalues_featurelist.sort_values()
pvalues_featurelist.head(10)
```

Out[4]:
```
id_number
F1039    4.028376e-13
F2201    3.386336e-11
F2955    4.471926e-11
F1417    1.217689e-10
F2675    2.118523e-10
F2784    2.630405e-10
F2663    8.330401e-10
F2930    1.497729e-09
F2874    2.037131e-09
F3182    2.161996e-09
dtype: float64
```

In [5]:
```
# how many features have p < 0.001
pvalues_featurelist[pvalues_featurelist<0.001].shape
```

Out[5]: (73,)

##### Summary

We got 73 features in this table that differ between 12C and 13C samples with p < 0.001.

We'll next look at their corresponding compound and intensity across ions and samples.

#### Part II. Pre-annotation and plot using khipu

In [6]:
```
!pip install --upgrade khipu-metabolomics
```

```
Collecting khipu-metabolomics
  Downloading https://files.pythonhosted.org/packages/11/b1/de56fe57c153d6009a9924b9a79f42e5a52cfb74928d8619eee3a6eb2
f9a/khipu_metabolomics-0.4.17-py3-none-any.whl
Requirement already satisfied, skipping upgrade: networkx in /opt/conda/lib/python3.7/site-packages (from khipu-metab
olomics) (2.3)
Requirement already satisfied, skipping upgrade: treelib in /opt/conda/lib/python3.7/site-packages (from khipu-metabo
lomics) (1.6.1)
Requirement already satisfied, skipping upgrade: mass2chem in /opt/conda/lib/python3.7/site-packages (from khipu-meta
bolomics) (0.4.1)
Requirement already satisfied, skipping upgrade: decorator>=4.3.0 in /opt/conda/lib/python3.7/site-packages (from net
workx->khipu-metabolomics) (4.4.0)
Requirement already satisfied, skipping upgrade: future in /opt/conda/lib/python3.7/site-packages (from treelib->khip
u-metabolomics) (0.18.2)
Requirement already satisfied, skipping upgrade: scipy in /opt/conda/lib/python3.7/site-packages (from mass2chem->khi
pu-metabolomics) (1.3.1)
Requirement already satisfied, skipping upgrade: numpy in /opt/conda/lib/python3.7/site-packages (from mass2chem->khi
pu-metabolomics) (1.17.2)
Installing collected packages: khipu-metabolomics
  Found existing installation: khipu-metabolomics 0.4.16
    Uninstalling khipu-metabolomics-0.4.16:
      Successfully uninstalled khipu-metabolomics-0.4.16
Successfully installed khipu-metabolomics-0.4.17
```

In [7]:
```
from khipu.extended import *
```

In [8]:
```
# One can redefine the patterns
# isotope_search_patterns = isotope_search_patterns[:6]

for x in (adduct_search_patterns, isotope_search_patterns, extended_adducts):
    print(x, '\n')
```

```
[(21.982, 'Na/H'), (41.026549, 'ACN'), (35.9767, 'HCl'), (37.955882, 'K/H')]

[(1.003355, '13C/12C', (0, 0.8)), (2.00671, '13C/12C*2', (0, 0.8)), (3.010065, '13C/12C*3', (0, 0.8)), (4.01342, '13C
/12C*4', (0, 0.8)), (5.016775, '13C/12C*5', (0, 0.8)), (6.02013, '13C/12C*6', (0, 0.8)), (7.023485, '13C/12C*7', (0,
0.8)), (8.02684, '13C/12C*8', (0, 0.8)), (9.030195, '13C/12C*9', (0, 0.8)), (10.03355, '13C/12C*10', (0, 0.8)), (11.0
36905, '13C/12C*11', (0, 0.8)), (12.04026, '13C/12C*12', (0, 0.8))]

[(1.0078, 'H'), (-1.0078, '-H'), (10.991, 'Na/H, double charged'), (0.5017, '13C/12C, double charged'), (117.02655, '
-NH3'), (17.02655, 'NH3'), (-18.0106, '-H2O'), (18.0106, 'H2O'), (18.033823, 'NH4'), (27.01089904, 'HCN'), (37.94694,
'Ca/H2'), (32.026215, 'MeOH'), (43.96389, 'Na2/H2'), (67.987424, 'NaCOOH'), (83.961361, 'KCOOH'), (97.96737927, 'H2SO
4'), (97.97689507, 'H3PO4')]
```

In [9]:
```
# read data via khipu
peaklist = read_features_from_text(open("ecoli_pos.tsv").read(),
                    id_col=0, mz_col=1, rtime_col=2, intensity_cols=(3, 9), delimiter="\t")
```

```
table header looks like:  ['id_number', 'mz', 'rtime', '12C_Ecoli_20220321_004', '12C_Ecoli_20220321_004_202203220950
30', '12C_Ecoli_20220321_004_20220322130235', '13C_Ecoli_20220321_004', '13C_Ecoli_20220321_004_20220322132355', '13C
_Ecoli_20220321_004_20220322101150']
Read 3602 feature lines
```

In [10]:
```
peaklist[55]
```

Out[10]:
```
{'id': 'F56',
 'mz': 97.1013,
 'rtime': 66.21,
 'intensities': [3138738.0,
  2184196.0,
  1433236.0,
  3032564.0,
  1719115.0,
  1937016.0],
 'representative_intensity': 2240810.8333333335}
```

In [11]:
```
khipu_list, all_assigned_peaks = peaklist_to_khipu_list(peaklist,
                    isotope_search_patterns=isotope_search_patterns,
                    adduct_search_patterns=adduct_search_patterns,
                    mz_tolerance_ppm=5,
                    rt_tolerance=2,
                    mode='pos')
```

```
Initial khipu search grid:
                M+H+       Na/H        HCl        K/H        ACN
M0           1.007276  22.989276  36.983976  38.963158  42.033825
13C/12C      2.010631  23.992631  37.987331  39.966513  43.037180
13C/12C*2    3.013986  24.995986  38.990686  40.969868  44.040535
13C/12C*3    4.017341  25.999341  39.994041  41.973223  45.043890
13C/12C*4    5.020696  27.002696  40.997396  42.976578  46.047245
13C/12C*5    6.024051  28.006051  42.000751  43.979933  47.050600
13C/12C*6    7.027406  29.009406  43.004106  44.983288  48.053955
13C/12C*7    8.030761  30.012761  44.007461  45.986643  49.057310
13C/12C*8    9.034116  31.016116  45.010816  46.989998  50.060665
13C/12C*9   10.037471  32.019471  46.014171  47.993353  51.064020
13C/12C*10  11.040826  33.022826  47.017526  48.996708  52.067375
13C/12C*11  12.044181  34.026181  48.020881  50.000063  53.070730
13C/12C*12  13.047536  35.029536  49.024236  51.003418  54.074085


Warning no match in isotope_pattern:  16.053999999999988
Unknown isotope match ~  (137.1188, 'F292')
Warning no match in isotope_pattern:  17.057299999999984
Unknown isotope match ~  (138.1221, 'F397')
Warning no match in isotope_pattern:  15.050899999999984
Unknown isotope match ~  (152.1378, 'F1251')
Warning no match in isotope_pattern:  16.05419999999998
Unknown isotope match ~  (153.1411, 'F1318')
Warning no match in isotope_pattern:  15.050899999999984
Unknown isotope match ~  (152.1378, 'F1252')
Warning no match in isotope_pattern:  16.05419999999998
Unknown isotope match ~  (153.1411, 'F1319')
Warning no match in isotope_pattern:  16.0539
Unknown isotope match ~  (123.1031, 'F570')
Warning no match in isotope_pattern:  13.0428
Unknown isotope match ~  (181.1392, 'F1967')
Warning no match in isotope_pattern:  17.05710000000002
Unknown isotope match ~  (194.1596, 'F1548')
Warning no match in isotope_pattern:  13.043700000000001
Unknown isotope match ~  (176.0708, 'F1657')
Warning no match in isotope_pattern:  16.053000000000026
Unknown isotope match ~  (259.0966, 'F3572')
Warning no match in isotope_pattern:  15.050000000000011
Unknown isotope match ~  (291.1191, 'F3517')
Warning no match in isotope_pattern:  26.086900000000014
Unknown isotope match ~  (302.156, 'F2566')
Warning no match in isotope_pattern:  37.122299999999996
Unknown isotope match ~  (313.1914, 'F3577')
Warning no match in isotope_pattern:  14.04480000000001
Unknown isotope match ~  (320.1205, 'F2769')
Warning no match in isotope_pattern:  13.041999999999973
Unknown isotope match ~  (347.0852, 'F3097')
Warning no match in isotope_pattern:  22.074899999999957
Unknown isotope match ~  (344.1819, 'F3403')
Warning no match in isotope_pattern:  19.06310000000002
Unknown isotope match ~  (294.1329, 'F2385')
```

In [12]:
```
len(khipu_list), len(all_assigned_peaks)
```

Out[12]: (548, 1746)

In [13]:
```
ksizes = [
    (len(KP.nodes_to_use), KP.id) for KP in khipu_list
]
ksizes = sorted(ksizes, reverse=True)
print(ksizes[:30])
```

```
[(30, 'kp31_114.0549'), (20, 'kp33_116.0479'), (16, 'kp90_103.0165'), (15, 'kp11_123.0222'), (13, 'kp132_187.1686'),
(11, 'kp259_309.0862'), (10, 'kp93_165.079'), (10, 'kp40_136.0998'), (10, 'kp198_231.1861'), (9, 'kp94_129.1024'), (9
, 'kp65_123.0318'), (9, 'kp5_122.084'), (9, 'kp2_71.0008'), (9, 'kp211_236.0834'), (8, 'kp201_174.1007'), (7, 'kp54_9
7.0985'), (7, 'kp357_233.125'), (7, 'kp284_414.2045'), (7, 'kp139_181.0741'), (7, 'kp135_112.0156'), (7, 'kp10_113.08
42'), (7, 'kp100_108.0213'), (6, 'kp309_333.0342'), (6, 'kp308_307.0841'), (6, 'kp29_97.968'), (6, 'kp291_283.8771'),
(6, 'kp266_305.0676'), (6, 'kp224_243.0859'), (6, 'kp177_187.8941'), (6, 'kp172_134.037')]
```

In [14]:
```
# build two dictionary to use.
# kpdict indexed by khipu ID, fdict indexed by feature ID
kpdict, fdict = {}, {}
for KP in khipu_list:
    kpdict[KP.id] = KP
    for f in KP.nodes_to_use:
        fdict[f] = KP
```

In [15]:
```
kpdict['kp11_123.0222'].plot()
```

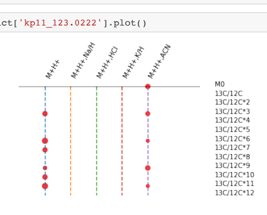


**Examine the most differential features we got above**

In [16]:
```
sig = list(pvalues_featurelist[pvalues_featurelist<0.001].index)
print(sig)
```

```
['F1039', 'F2201', 'F2955', 'F1417', 'F2675', 'F2784', 'F2663', 'F2930', 'F2874', 'F3182', 'F2178', 'F2633', 'F3024',
'F2931', 'F2338', 'F2923', 'F2863', 'F2132', 'F2979', 'F3078', 'F2343', 'F3009', 'F2421', 'F3042', 'F2924', 'F2706',
'F2727', 'F2653', 'F2514', 'F2716', 'F3181', 'F3180', 'F2720', 'F2695', 'F1843', 'F1703', 'F1691', 'F2422', 'F2667',
'F780', 'F3054', 'F1177', 'F2785', 'F2707', 'F2246', 'F2203', 'F2544', 'F894', 'F1833', 'F2303', 'F1191', 'F2946', 'F
3103', 'F2711', 'F3031', 'F3364', 'F2987', 'F2247', 'F65', 'F1495', 'F1690', 'F2950', 'F1496', 'F1894', 'F499', 'F302
2', 'F2562', 'F390', 'F3362', 'F1508', 'F2572', 'F3102', 'F3494']
```

In [17]:
```
# get khipu IDs for the above features
sig_kps = set([])
for x in sig:
    if x in fdict:
        sig_kps.add( fdict[x].id )

print(len(sig_kps), sig_kps)
```

```
25 {'kp232_242.0807', 'kp304_577.5973', 'kp325_206.0694', 'kp259_309.0862', 'kp357_233.125', 'kp216_183.0898', 'kp224
_243.0859', 'kp266_305.0676', 'kp366_327.0911', 'kp302_486.3565', 'kp391_365.0897', 'kp371_244.9836', 'kp263_306.5779
', 'kp303_577.0978', 'kp240_297.0904', 'kp537_326.0325', 'kp167_156.079', 'kp276_401.0192', 'kp111_146.0691', 'kp469_
91.5987', 'kp248_260.1324', 'kp353_202.0414', 'kp279_132.0899', 'kp246_219.9739', 'kp433_419.05'}
```

In [18]:
```
# show some sig_kps
sig_kps = list(sig_kps)
for i in sig_kps[:5]:
    print(kpdict[i].id, kpdict[i].neutral_mass)
    print(kpdict[i].khipu_grid)
    print('\n')
```

```
kp232_242.0807 242.08066853322998
             M+H+
M0          F3330
13C/12C*11  F2343
13C/12C*12  F2355


kp304_577.5973 577.59734603323
          M+H+ M+H+,13C/12C, double charged
M0       F3010
13C/12C  F3022                         F3024


kp325_206.0694 206.06939853323
             M+H+
M0          F3191
13C/12C*10  F1417


kp259_309.0862 309.0862428189443
             M+H+   Na/H HCl K/H ACN
M0                 F3324
13C/12C
13C/12C*2   F2695
13C/12C*3
13C/12C*4
13C/12C*5
13C/12C*6   F2729
13C/12C*7   F2740
13C/12C*8
13C/12C*9   F3578  F3004
13C/12C*10         F3050
13C/12C*11
13C/12C*12


kp357_233.125 233.12496553351826
             M+H+ Na/H    HCl    K/H ACN HCl,NH3
M0                       F3429              F3574
13C/12C
13C/12C*2
13C/12C*3
13C/12C*4
13C/12C*5
13C/12C*6
13C/12C*7
13C/12C*8
13C/12C*9
13C/12C*10  F3332        F3147  F3181
13C/12C*11
13C/12C*12
```

In [19]:
```
# closer look
kpdict['kp246_219.9739'].khipu_grid
```

Out[19]:

| | M+H+ | Na/H | HCl | K/H | ACN | M+H+,-H2O |
|---|---|---|---|---|---|---|
| M0 | F1483 | | | F2496 | | F2092 |
| 13C/12C | | | | | | |
| 13C/12C*2 | | | | | | |
| 13C/12C*3 | | | | | | |
| 13C/12C*4 | | | | | | |
| 13C/12C*5 | | | | F3362 | | |
| 13C/12C*6 | | | | | | |
| 13C/12C*7 | | | | | | |
| 13C/12C*8 | | | | | | |
| 13C/12C*9 | | | | | | |
| 13C/12C*10 | | | | | | |
| 13C/12C*11 | | | | | | |
| 13C/12C*12 | | | | | | |

In [20]:
```
# m/z view
kpdict['kp246_219.9739'].print3()
```

Out[20]:

| | M+H+ | Na/H | HCl | K/H | ACN | M+H+,-H2O |
|---|---|---|---|---|---|---|
| M0 | 220.982 | | | 258.938 | | 202.971 |
| 13C/12C | | | | | | |
| 13C/12C*2 | | | | | | |
| 13C/12C*3 | | | | | | |
| 13C/12C*4 | | | | | | |
| 13C/12C*5 | | | | 263.953 | | |
| 13C/12C*6 | | | | | | |
| 13C/12C*7 | | | | | | |
| 13C/12C*8 | | | | | | |
| 13C/12C*9 | | | | | | |
| 13C/12C*10 | | | | | | |
| 13C/12C*11 | | | | | | |
| 13C/12C*12 | | | | | | |

In [21]:
```
for i in sig_kps[10: ]:
    print(kpdict[i].id, kpdict[i].neutral_mass)
    print(kpdict[i].khipu_grid)
    print('\n')
```

```
kp391_365.0897 365.08974353323
             M+H+
M0          F3406
13C/12C*12  F2675


kp371_244.9836 244.98358603322998
            M+H+
M0         F3334
13C/12C*5  F2246


kp263_306.5779 306.57791853323
            M+H+
M0         F3567
13C/12C*7  F2716
13C/12C*8  F2722
13C/12C*9  F2732


kp303_577.0978 577.09779603323
          M+H+
M0       F3009
13C/12C  F3019


kp240_297.0904 297.09037103323
             M+H+
M0          F2454
13C/12C*11  F2663


kp537_326.0325 326.03247353323
    M+H+   Na/H M+H+,NH3
M0  F2867  F3103    F3064


kp167_156.079 156.07898886656332
             M+H+ Na/H HCl K/H    ACN
M0          F1628                F1775
13C/12C     F1690
13C/12C*2
13C/12C*3
13C/12C*4
13C/12C*5
13C/12C*6
13C/12C*7
13C/12C*8
13C/12C*9
13C/12C*10
13C/12C*11
13C/12C*12


kp276_401.0192 401.01917873323
             M+H+ Na/H HCl    K/H ACN
M0                           F3552
13C/12C     F3504
13C/12C*2
13C/12C*3
13C/12C*4
13C/12C*5
13C/12C*6
13C/12C*7
13C/12C*8
13C/12C*9
13C/12C*10  F2784            F2884
13C/12C*11  F2789
13C/12C*12


kp111_146.0691 146.06913353323
            M+H+
M0         F2896
13C/12C*3  F1092
13C/12C*4  F1177
13C/12C*5  F1239


kp469_91.5987 91.59870720009425
    M+H+    HCl   ACN M+H+,H2O HCl,-H2O M+H+,H3PO4
M0  F356  F1508  F468     F336     F320      F1423


kp248_260.1324 260.13244603323
          M+H+
M0       F2514
13C/12C  F2522


kp353_202.0414 202.04138603322997
            M+H+
M0         F3285
13C/12C*5  F2203


kp279_132.0899 132.08993603322998
            M+H+
M0         F2795
13C/12C*5   F390


kp246_219.9739 219.97391053323
             M+H+ Na/H HCl    K/H ACN M+H+,-H2O
M0          F1483            F2496         F2092
13C/12C
13C/12C*2
13C/12C*3
13C/12C*4
13C/12C*5                    F3362
13C/12C*6
13C/12C*7
13C/12C*8
13C/12C*9
13C/12C*10
13C/12C*11
13C/12C*12


kp433_419.05 419.05004853323
             M+H+
M0          F3536
13C/12C*10  F2863
```

In [22]:
```
kpdict['kp224_243.0859'].format_to_epds()
```

Out[22]:
```
{'interim_id': 'root@243.0859',
 'neutral_formula_mass': 243.0858820335399,
 'neutral_formula': None,
 'Database_referred': [],
 'identity': [],
 'MS1_pseudo_Spectra': [{'id': 'F3331',
   'mz': 244.093,
   'rtime': 27.05,
   'intensities': [3324402.0, 4072782.0, 3961633.0, 0.0, 0.0, 0.0],
   'representative_intensity': 1893136.1666666667,
   'parent_masstrack_id': '244.093',
   'isotope': 'M0',
   'modification': 'M+H+',
   'ion_relation': 'M0,M+H+'},
  {'id': 'F3472',
   'mz': 291.0793,
   'rtime': 27.27,
   'intensities': [0.0, 0.0, 0.0, 553698.0, 1014278.0, 1075498.0],
   'representative_intensity': 440579.0,
   'parent_masstrack_id': '291.0793',
   'isotope': '13C/12C*9',
   'modification': 'K/H',
   'ion_relation': '13C/12C*9,K/H'},
  {'id': 'F2318',
   'mz': 253.1233,
   'rtime': 27.5,
   'intensities': [0.0, 0.0, 0.0, 3996975.0, 4208279.0, 4402969.0],
   'representative_intensity': 2101370.5,
   'parent_masstrack_id': '253.1233',
   'isotope': '13C/12C*9',
   'modification': 'M+H+',
   'ion_relation': '13C/12C*9,M+H+'},
  {'id': 'F3483',
   'mz': 282.0491,
   'rtime': 27.27,
   'intensities': [495468.0, 1015145.0, 1119976.0, 0.0, 0.0, 0.0],
   'representative_intensity': 438431.5,
   'parent_masstrack_id': '282.0491',
   'isotope': 'M0',
   'modification': 'K/H',
   'ion_relation': 'M0,K/H'},
  {'id': 'F2158',
   'mz': 245.0967,
   'rtime': 27.73,
   'intensities': [420479.0, 355600.0, 241481.0, 96994.0, 0.0, 55216.0],
   'representative_intensity': 194961.66666666666,
   'parent_masstrack_id': '245.0967',
   'isotope': '13C/12C',
   'modification': 'M+H+',
   'ion_relation': '13C/12C,M+H+'},
  {'id': 'F2303',
   'mz': 252.1199,
   'rtime': 27.27,
   'intensities': [0.0, 0.0, 0.0, 0.0, 526192.0, 744605.0],
   'representative_intensity': 211799.5,
   'parent_masstrack_id': '252.1199',
   'isotope': '13C/12C*8',
   'modification': 'M+H+',
   'ion_relation': '13C/12C*8,M+H+'}],
 'MS2_Spectra': []}
```

**We can extract sample data from khipu and plot in different ways**

In [23]:
```
kpdict['kp224_243.0859'].plot()
```

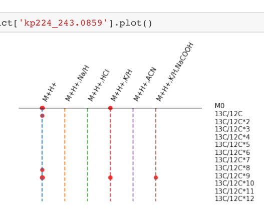


**Bar plot of average intensity per isotope in M+H+**

In [24]:
```
kpdict['kp224_243.0859'].get_khipu_intensities()['M+H+'].plot.bar()
```

Out[24]: <matplotlib.axes._subplots.AxesSubplot at 0x7f3d6cb63fd0>

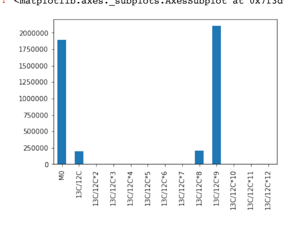


**Bar plot of all 6 samples**

In [25]:
```
sample_labels = ['12C', '12C', '12C', '13C', '13C', '13C']
```

In [26]:
```
def get_sample_data(kp, ion, sample_labels):
    '''kp as khipu instance. Assume sample_labels are in the same order as columns read into khipu instance.
    Return dataframe'''
    data = []
    for ii in range(len(sample_labels)):
        tmp = []
        for f in kp.khipu_grid[ion].values:
            if f:
                tmp.append(kp.feature_dict[f]['intensities'][ii])
            else:
                tmp.append(0)
        data.append(tmp)

    return pd.DataFrame(data,
                        columns=kp.khipu_grid.index,
                        index=sample_labels,
                        ).T
```

In [27]:
```
data = get_sample_data(kpdict['kp224_243.0859'], 'M+H+', sample_labels)
data
```

Out[27]:

| | 12C | 12C | 12C | 13C | 13C | 13C |
|---|---|---|---|---|---|---|
| M0 | 3324402.0 | 4072782.0 | 3961633.0 | 0.0 | 0.0 | 0.0 |
| 13C/12C | 420479.0 | 355600.0 | 241481.0 | 96994.0 | 0.0 | 55216.0 |
| 13C/12C*2 | 0.0 | 0.0 | 0.0 | 0.0 | 0.0 | 0.0 |
| 13C/12C*3 | 0.0 | 0.0 | 0.0 | 0.0 | 0.0 | 0.0 |
| 13C/12C*4 | 0.0 | 0.0 | 0.0 | 0.0 | 0.0 | 0.0 |
| 13C/12C*5 | 0.0 | 0.0 | 0.0 | 0.0 | 0.0 | 0.0 |
| 13C/12C*6 | 0.0 | 0.0 | 0.0 | 0.0 | 0.0 | 0.0 |
| 13C/12C*7 | 0.0 | 0.0 | 0.0 | 0.0 | 0.0 | 0.0 |
| 13C/12C*8 | 0.0 | 0.0 | 0.0 | 0.0 | 526192.0 | 744605.0 |
| 13C/12C*9 | 0.0 | 0.0 | 0.0 | 3996975.0 | 4208279.0 | 4402969.0 |
| 13C/12C*10 | 0.0 | 0.0 | 0.0 | 0.0 | 0.0 | 0.0 |
| 13C/12C*11 | 0.0 | 0.0 | 0.0 | 0.0 | 0.0 | 0.0 |
| 13C/12C*12 | 0.0 | 0.0 | 0.0 | 0.0 | 0.0 | 0.0 |

In [28]:
```
data.plot(kind='bar', stacked=False, figsize=(10, 4))
```

Out[28]: <matplotlib.axes._subplots.AxesSubplot at 0x7f3d6cc2c4e0>

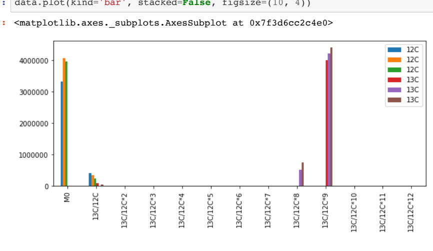


**The Acetylspermidine example**

In [29]:
```
fdict['F2353'].khipu_grid
```

Out[29]:

| | M+H+ | Na/H | HCl | K/H | ACN | M+H+,Na2/H2 |
|---|---|---|---|---|---|---|
| M0 | F2353 | F3209 | | F3269 | | |
| 13C/12C | F1331 | | | | | |
| 13C/12C*2 | F1397 | | | | | |
| 13C/12C*3 | F1434 | | | | | |
| 13C/12C*4 | F1461 | | | | | |
| 13C/12C*5 | F1507 | | | | | |
| 13C/12C*6 | F1550 | | | | | |
| 13C/12C*7 | | | | | | |
| 13C/12C*8 | F1685 | | | | | F2000 |
| 13C/12C*9 | F1741 | F1454 | | F1883 | | |
| 13C/12C*10 | | | | | | |
| 13C/12C*11 | | | | | | |
| 13C/12C*12 | | | | | | |

In [30]:
```
fdict['F2353'].plot()
```

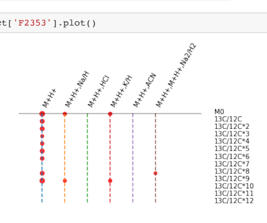


In [31]:
```
KP2353=fdict['F2353']
data = get_sample_data(KP2353, 'M+H+', sample_labels)
fig = data.plot(kind='bar', stacked=False, width=0.7, figsize=(10, 4))
f = fig.get_figure()
f.tight_layout()
f.savefig('KP2353_samples.pdf')
```

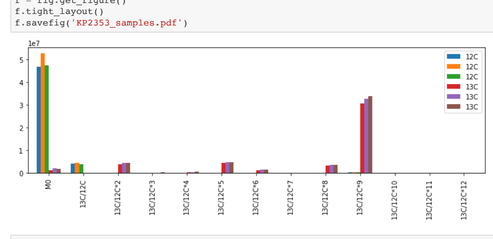


In [32]:
```
KP2353.neutral_mass
# HMDB matches to N1 or N8- Acetylspermidine    C9H21N3O        187.1685
```

Out[32]: 187.16862014861462

In [35]:
```
# options to save PDF and replace trunk labels
# one can define own relabel_dict
plot_khipugram(KP2353.get_khipu_intensities(), savepdf='khipugram_F2353.pdf', relabel=relabel_dict)
```

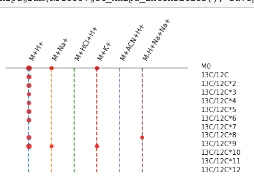


#### Part III. Data formats in khipu and interface with other tools

- khipu exports pre-annotation to two files of JSON and tab delimited formats, this_test.json and this_test.tsv.
- Internal functions and data frames are available to import and use

In [34]:
```
# check what objects were imported
print([x for x in dir() if x[0]!='_'])
```

```
['In', 'KP', 'KP2353', 'Khipu', 'Out', 'PROTON', 'Weavor', 'add_data_to_tag', 'adduct_search_patterns', 'adduct_searc
h_patterns_neg', 'all_assigned_peaks', 'assign_masstrack_ids_in_khipu', 'build_centurion_tree', 'curve_fit', 'data',
'ecoli', 'electron', 'exit', 'export_empCpd_khipu_list', 'export_json_khipu_list', 'export_json_trees', 'export_khipu
_table', 'export_tsv_trees', 'extend_khipu_list', 'extended_adducts', 'f', 'fdict', 'fig', 'find_all_matches_centurio
n_indexed_list', 'find_trees_by_datatag', 'find_trees_by_datatag_list', 'get_adduct_edge_pairs', 'get_ipython', 'get_
isotope_pattern_name', 'get_isotopic_edge_pairs', 'get_sample_data', 'graphs_to_khipu_list', 'i', 'is_datatag_in_tree
', 'isotope_search_patterns', 'json', 'khipu_annotate', 'khipu_diagnosis', 'khipu_list', 'kpdict', 'ksizes', 'local_r
ead_file', 'make_edge_tag', 'make_expected_adduct_index', 'make_peak_dict', 'make_peak_tag', 'np', 'nx', 'pd', 'peakl
ist', 'peaklist_to_khipu_list', 'peaks_to_networks', 'plot_khipugram', 'plt', 'pvalues_featurelist', 'quit', 'read_fe
atures_from_text', 'realign_isotopes', 'realign_isotopes_reverse', 'relabel_dict', 'rt_compared_by_values', 'rt_match
ed_by_tolerance', 'sample_labels', 'sig', 'sig_kps', 'treelib', 'ttest', 'ttest_ind', 'x']
```

In [36]:
```
# get documentation on the khipu class
help(KP)
```

```
Help on Khipu in module khipu.model object:

class Khipu(builtins.object)
 |  Khipu(subnetwork)
 |
 |  2-tier tree representation of an empirical compound.
 |  An empTree follows the matrix notion of a family of peaks: a tree of 2 levels - isotopes and adducts.
 |  A true root of khipu is the compound in neutral formula.
 |  A peudo root of khipu is the ion of lowest m/z.
```

```
 |  The representative adduct should have the most abundance.
 |  The matrix notion of a khipu is a DataFrame: columns are Adduct labels; rows isotopes.
 |  trunk : default for adducts
 |  branch : default for isotopes
 |
 |  Methods defined here:
 |
 |  __init__(self, subnetwork)
 |      Do not include negative m/z difference in this class, as the node of lowest m/z is used as root.
 |      Neutral loss and fragments can be searched after khipu is established.
 |      Input subnetwork is undirected graph, but khipu will require directed graph/tree, by increasing m/z.
 |      subnetwork : undirected graph. Example edges:
 |          ('F3533', 'F20', {'type': 'modification', 'tag': 'Na/H'}),
 |          ('F195', 'F20', {'type': 'modification', 'tag': 'Acetonitrile'}),
 |          ('F20', 'F53', {'type': 'isotope', 'tag': '13C/12C'}),
 |
 |  branch_abstraction(self, isotopic_edges, adduct_edges)
 |      Abstract a group of connecgted isotopic featrures into a branch.
 |      Reduce the input network to a set of abstracted_adduct_edges (hence new tree) of B-nodes.
 |
 |      Returns
 |      -------
 |      abstracted_adduct_edges : list of nonredundant directed edges with data tag
 |      branch_dict : dictionary, branch ID to member features/nodes.
 |
 |      Notes
 |      -----
 |      Membership of B-nodes is returned as branch_dict, which is needed to realign to khipu grid.
 |      Without branch constraint, the grid realignment is error prone.
 |      Not checking if abstracted_adduct_edges are fully connected.
 |
 |  build_khipu(self, WeavorInstance, mz_tolerance_ppm=5, check_fitness=False)
 |      Convert a network of two types of edges, isotopic and adduct, to khipu instances of unique nodes.
 |      Use the grid to enforce unique ion in each position, as the initial network can contain
 |      erroneously redundant leaves/nodes.
 |
 |      WeavorInstance : Weavor instance to solve the grid.
 |      check_fitness : if True, chech if replacing with any redundant nodes improves fitness.
 |          This is a placeholder as a good fitness function is not easy to implement and it slows down computing.
 |          Future use can be if check_fitness: self.select_by_fitness()
 |
 |      Updates
 |      -------
 |      self.khipu_grid : pd.DataFrame
 |      self.neutral_mass : inferred neutral mass for the khipu compound
 |
 |  clean(self, WeavorInstance, mz_tolerance_ppm)
 |      Clean up the input subnetwork, only using unique features to build a khipu frame.
 |      Redundant features are kept aside. The leftover features will be sent off to build new khipus.
 |
 |      Updates
 |      -------
 |      self.feature_dict
 |      self.mzstr_dict
 |      self.median_rtime
 |      self.nodes_to_use
 |      self.redundant_nodes
 |      self.sorted_mz_peak_ids
 |      self.root       # temporary
 |
 |      Notes
 |      -----
 |      Note: mzstr_dict can be problematic in data that were not processed well,
 |      because minor potential shift can confuse ion relations.
 |      This clean() may cut some initial edges.
 |
 |  down_size(self)
 |      If input_network is too big, down size by selected the features of highest abundance.
 |      The extra nodes are not kept, because they can be picked up by extended search later if they fit this Khipu.
 |
 |  export_json(self)
 |      Placeholder.
 |
 |  export_json_grid_data(self)
 |
 |  extended_search(self, mztree, adduct_search_patterns_extended, mz_tolerance_ppm=5, rt_tolerance=2)
 |      Find additional adducts from unassigned_peaks using adduct_search_patterns_extended.
 |      mztree here are indexed unassigned_peaks.
 |
 |      Updates
 |      -------
 |      self.feature_map
 |      self.khipu_grid
 |
 |      Returns
 |      -------
 |      added_peaks
 |
 |      Notes
 |      -----
 |      annotation_dict may have fewer nodes than nodes_to_use, but is preferred here for cleaner results.
 |
 |  format_to_epds(self, id='')
 |      Format khipu to empirical compound, with added ion notions.
 |      A small number of features do not map the khipu grid (e.g. their edges violate DAG rules).
 |      They are kept in empCpd as "undetermined".
 |
 |  get_feature_dict(self, peak_dict, mz_tolerance_ppm)
 |      Index all input features; establish str identifier for features of same/close m/z values.
 |      Base on asari mass track IDs; keep unique m/z only.
 |      It's more efficient to use, since feature_dict is much smaller than peak_dict.
 |
 |      Parameters
 |      ----------
 |      peak_dict : dict of peaks/features indexed by IDs. Must have fields
 |                  'id', 'mz', 'rtime', 'representative_intensity'.
 |      mz_tolerance_ppm : ppm tolerance in examining m/z groups.
 |
 |      Returns
 |      -------
 |      feature_dict : feature_dict with 'parent_masstrack_id'.
 |      mzstr_dict : dict indexed by parent_masstrack_id, e.g. {'trackx': [f1, f2], ...}
 |
 |  get_khipu_intensities(self)
 |      Return abundance_matrix as DataFrame in same layout as self.khipu_grid
 |
 |  get_khipu_mzgrid_print(self)
 |      Return str m/z matrix as DataFrame in same layout as self.khipu_grid, for visual purpose.
 |
 |  get_pruned_network(self)
 |      Get extra features and edges that are not fit in this khipu.
 |      Updates:
 |      self.redundant_nodes
 |      self.pruned_network
 |
 |  plot = plot_khipu_diagram(self, savepdf='')
 |
 |  plot2 = plot_khipu_diagram_rotated(self)
 |
 |  plot_khipu_diagram(self, savepdf='')
 |      Plot the khipu grid as diagram.
 |      Use MatPlotLib as default engine.
 |
 |  plot_khipu_diagram_rotated(self)
 |
 |  print = print_khipu(self)
 |
 |  print2 = print_khipu_rotated(self)
 |
 |  print3 = get_khipu_mzgrid_print(self)
 |
 |  print_khipu(self)
 |      Print khipu using adducts as trunk and isotopes as branches (preferred)
 |
 |  print_khipu_rotated(self)
 |      Print khipu using isotopes as trunk and adducts as branches
 |
 |  sort_nodes_by_mz(self)
 |      sort the nodes by increasing m/z
 |
 |  ----------------------------------------------------------------------
 |  Data descriptors defined here:
 |
 |  __dict__
 |      dictionary for instance variables (if defined)
 |
 |  __weakref__
 |      list of weak references to the object (if defined)
```

In [37]:

```
# get documentation on a function, e.g. peaks_to_networks
help(peaks_to_networks)
```

```
Help on function peaks_to_networks in module khipu.utils:

peaks_to_networks(peak_list, isotope_search_patterns=[(1.003355, '13C/12C', (0, 0.8)), (2.00671, '13C/12C*2', (0, 0.8
)), (3.010065, '13C/12C*3', (0, 0.8)), (4.01342, '13C/12C*4', (0, 0.8)), (5.016775, '13C/12C*5', (0, 0.8)), (6.02013,
'13C/12C*6', (0, 0.8))], adduct_search_patterns=[(21.982, 'Na/H'), (41.026549, 'ACN')], mz_tolerance_ppm=5, rt_tolera
nce=2)
    Search peak_list for patterns of isotopes and adducts, form a network and get connected subnetworks.

    Parameters
    ----------
    list_peaks : [{'mz': 133.09702315984987, 'rtime': 654, 'id': 555}, ...]

    isotope_search_patterns : exact list used to retrieve the subnetworks. E.g.
        [ (1.003355, '13C/12C', (0, 0.8)),
        (2.00671, '13C/12C*2', (0, 0.8)),
        (3.010065, '13C/12C*3', (0, 0.8)),
        (4.01342, '13C/12C*4', (0, 0.8)),
        (5.016775, '13C/12C*5', (0, 0.8)),
        (6.02013, '13C/12C*6', (0, 0.8)),]

    adduct_search_patterns : exact list used to retrieve the subnetworks.
        It's not recommended to have a long list here, as it's better to search additional
        in-source modifications after empCpds are seeded. Example adduct_search_patterns list:
        [ (1.0078, 'H'),
        (21.9820, 'Na/H'),
        (41.026549, 'Acetonitrile')]

    mz_tolerance_ppm : ppm tolerance in examining m/z patterns.
    rt_tolerance: tolerance threshold for deviation in retetion time, arbitrary unit depending on input data.
            Default intended as 2 seconds.

    Returns
    -------
    subnetwork : undirected graph. Example edges:
        [('F1606', 'F20', {'type': 'modification', 'tag': 'H'}),
        ('F3533', 'F20', {'type': 'modification', 'tag': 'Na/H'}),
        ('F195', 'F20', {'type': 'modification', 'tag': 'Acetonitrile'}),
        ('F20', 'F807', {'type': 'modification', 'tag': 'Acetonitrile'}),
        ('F20', 'F53', {'type': 'isotope', 'tag': '13C/12C'}),
        ('F874', 'F808', {'type': 'isotope', 'tag': '13C/12C'})]

    peak_dict : JSON peaks indexed by ID
    edge_dict : edge_tag is str sorted, but the dict values preserve the direction, which is missed in nx.subnetwork.

    Notes
    -----
    Features of low abundance may not have detectable isotopes, but can have multiple adducts.
    Do not include too many adducts in the intial search.
    Do not include neutral loss and fragments in initial search.
    They are better done after a list of khipus are constructed.
```

In [38]:

```
# We have seen the JSON format earlier
fdict['F2353'].format_to_epds()
```

Out[38]:


```
{'interim_id': 'root@187.1686',
 'neutral_formula_mass': 187.16862014861462,
 'neutral_formula': None,
 'Database_referred': [],
 'identity': [],
 'MS1_pseudo_Spectra': [{'id': 'F1434',
   'mz': 191.1859,
   'rtime': 20.57,
   'intensities': [0.0, 0.0, 0.0, 256261.0, 179058.0, 284038.0],
   'representative_intensity': 119892.83333333333,
   'parent_masstrack_id': '191.1859',
   'isotope': '13C/12C*3',
   'modification': 'M+H+',
   'ion_relation': '13C/12C*3,M+H+'},
  {'id': 'F1507',
   'mz': 193.1927,
   'rtime': 20.81,
   'intensities': [0.0, 0.0, 0.0, 4399053.0, 4835021.0, 4901675.0],
   'representative_intensity': 2355958.1666666665,
   'parent_masstrack_id': '193.1927',
   'isotope': '13C/12C*5',
   'modification': 'M+H+',
   'ion_relation': '13C/12C*5,M+H+'},
  {'id': 'F3269',
   'mz': 226.1318,
   'rtime': 21.54,
   'intensities': [365144.0, 703817.0, 602973.0, 0.0, 0.0, 0.0],
   'representative_intensity': 278655.6666666667,
   'parent_masstrack_id': '226.1318',
   'isotope': 'M0',
   'modification': 'K/H',
   'ion_relation': 'M0,K/H'},
  {'id': 'F1883',
   'mz': 235.1621,
   'rtime': 21.54,
   'intensities': [0.0, 0.0, 0.0, 322390.0, 402996.0, 506915.0],
   'representative_intensity': 205383.5,
   'parent_masstrack_id': '235.1621',
   'isotope': '13C/12C*9',
   'modification': 'K/H',
   'ion_relation': '13C/12C*9,K/H'},
  {'id': 'F2353',
   'mz': 188.1759,
   'rtime': 20.57,
   'intensities': [46695047.0,
    52447244.0,
    47289802.0,
    1304564.0,
    2263739.0,
    1796286.0],
   'representative_intensity': 25299447.0,
   'parent_masstrack_id': '188.1759',
   'isotope': 'M0',
   'modification': 'M+H+',
   'ion_relation': 'M0,M+H+'},
  {'id': 'F3209',
   'mz': 210.1578,
   'rtime': 20.57,
   'intensities': [245212.0, 381771.0, 355978.0, 0.0, 0.0, 0.0],
   'representative_intensity': 163826.83333333334,
   'parent_masstrack_id': '210.1578',
   'isotope': 'M0',
   'modification': 'Na/H',
   'ion_relation': 'M0,Na/H'},
  {'id': 'F1461',
   'mz': 192.1893,
   'rtime': 20.81,
   'intensities': [0.0, 0.0, 0.0, 546925.0, 553011.0, 615354.0],
   'representative_intensity': 285881.6666666667,
   'parent_masstrack_id': '192.1893',
   'isotope': '13C/12C*4',
   'modification': 'M+H+',
   'ion_relation': '13C/12C*4,M+H+'},
  {'id': 'F1741',
   'mz': 197.2061,
   'rtime': 20.57,
   'intensities': [355590.0,
    430104.0,
    389484.0,
    30709282.0,
    32554138.0,
    33936088.0],
   'representative_intensity': 16395781.0,
   'parent_masstrack_id': '197.2061',
   'isotope': '13C/12C*9',
   'modification': 'M+H+',
   'ion_relation': '13C/12C*9,M+H+'},
  {'id': 'F1454',
   'mz': 219.1881,
   'rtime': 21.29,
   'intensities': [0.0, 0.0, 0.0, 111302.0, 228146.0, 227890.0],
   'representative_intensity': 94556.33333333333,
   'parent_masstrack_id': '219.1881',
   'isotope': '13C/12C*9',
   'modification': 'Na/H',
   'ion_relation': '13C/12C*9,Na/H'},
  {'id': 'F1331',
   'mz': 189.1792,
   'rtime': 20.57,
   'intensities': [4219502.0,
    4510375.0,
    4073666.0,
    138413.0,
    242879.0,
    191906.0],
   'representative_intensity': 2229456.8333333335,
   'parent_masstrack_id': '189.1792',
   'isotope': '13C/12C',
   'modification': 'M+H+',
   'ion_relation': '13C/12C,M+H+'},
  {'id': 'F1550',
   'mz': 194.196,
   'rtime': 20.81,
   'intensities': [0.0, 0.0, 0.0, 1356640.0, 1545713.0, 1555564.0],
   'representative_intensity': 742986.1666666666,
   'parent_masstrack_id': '194.196',
   'isotope': '13C/12C*6',
   'modification': 'M+H+',
   'ion_relation': '13C/12C*6,M+H+'},
  {'id': 'F1397',
   'mz': 190.1826,
   'rtime': 20.57,
   'intensities': [109712.0,
    153168.0,
    100640.0,
    4015179.0,
    4406835.0,
    4502197.0],
   'representative_intensity': 2214621.8333333335,
   'parent_masstrack_id': '190.1826',
   'isotope': '13C/12C*2',
   'modification': 'M+H+',
   'ion_relation': '13C/12C*2,M+H+'},
  {'id': 'F1685',
   'mz': 196.2028,
   'rtime': 20.81,
   'intensities': [0.0, 0.0, 0.0, 3403618.0, 3619416.0, 3676605.0],
   'representative_intensity': 1783273.1666666667,
   'parent_masstrack_id': '196.2028',
   'isotope': '13C/12C*8',
   'modification': 'M+H+',
   'ion_relation': '13C/12C*8,M+H+'}],
 'MS2_Spectra': []}
```


#### Summary

Pre-annotation by khipu allow us to use standard data science tools on isotope tracing metabolomics data.

User can further filter for preferred isotope patterns and apply custom models.
